# The Hypothesis Race Model for evaluation of research findings

**DOI:** 10.64898/2026.05.28.728385

**Authors:** Robert E. Kelly

## Abstract

Null Hypothesis Significance Testing (NHST) remains the dominant paradigm for evaluation of empirical research findings in medicine and the social sciences despite concerns about frequent misinterpretations of those findings. Achievement of “statistical significance,” the goal of NHST, often beckons unrealistic conclusions. Helpful would be the addition of a broader, Bayesian perspective of research in terms of progressive readjustment of hypothesis credibility from all sources of evidence. For this purpose, the Hypothesis Race Model (HRM) provides an intuitive Bayesian approach that builds upon NHST-concepts, helping to correct misunderstandings with minimal reeducation. The HRM is an extension of the Bayesian approach by Ioannidis in 2005 that helped to explain “why most published research findings are false.” It is powerful enough to serve as the foundation for mathematical models to estimate and reduce the cost of empirical hypothesis testing.

## 1. Introduction

Null hypothesis significance testing (NHST) has provided the backbone of our evidence-based medicine, but a statistically significant result alone does not answer the important question of how confident we can be that a given hypothesis is true [1–4]. Experience shows that results of such hypothesis testing often cannot be replicated [5–9], a problem known as the “replication crisis” [10–14]. Using common-sense assumptions and mathematics, John Ioannidis demonstrated why we might expect that most published research findings are false [15]; and subsequent testing has shown high rates of false-positive findings across multiple fields [16–22].

Our aims for hypothesis testing should be to

1. Reliably confirm true hypotheses and disconfirm false ones,
2. as cost-effectively as possible.

Our approach needs to work for all hypotheses tested. Any approach that in some cases tends to confirm or disconfirm hypotheses regardless of their truth value would defeat the purpose of hypothesis testing.

Bayesian approaches in principle hold the potential to accomplish these aims [1,3,23–25], but their use has been criticized for being too complex, beyond the educational training of most researchers, and for allowing too much freedom in assignment of model parameters, thereby allowing researchers to arbitrarily support conclusions of their choice, rather than conclusions dictated by empirical evidence [26–30]. Bayesian approaches necessarily make prior assumptions concerning the effect size (or its probability distribution) for the hypothesis tested, which may poorly approximate the actual effect size [31–33]. In addition, attempts to ascertain statistical significance from single empirical trials using a Bayesian approach do not differ meaningfully from those derived from standard NHST, apart from perhaps lowering the nominal *α*-level for statistical significance [34].

The prospect of applying Bayesian statistics to draw meaningful conclusions can seem daunting, even for the simplest “binary” application, where all test results are considered either positive or negative, statistically significant or nonsignificant. Uncertainty in conclusions stems not only from the stochastic nature of the testing process, but also from three of the model parameters, which cannot be known precisely: *R*_0_, *α*, and *β. R*_0_ is the estimated odds (probability true/probability false) that the hypothesis is true, prior to testing; *α* is the probability of a positive test given that the hypothesis is false (*α*-level, type I error for NHST); and *β* is the probability of a negative test given that the hypothesis is true (1 – power, type II error) [14,35]. Although the model value, *α*_m_, can generally be calculated precisely, the actual *α* is affected by multiple possible sources of bias that include the risk of methodological errors, publication bias, *p*-hacking, and research bias of all kinds (observer effect, placebo effect, selection bias, etc.) [36–43]. Finally, calculation of *β* and power require knowledge of the effect size corresponding to a true hypothesis [44], which cannot be known because determining whether the effect size is approximately zero is the purpose of the hypothesis testing.

Here I demonstrate that it is possible to navigate the uncertainties involved with applying Bayesian statistics to allow confirmation or disconfirmation of study hypotheses, while minimizing research costs. First, a Hypothesis Race Model (HRM) is described that provides a Bayesian framework with which to interpret research findings. The model can be utilized as an extension of Null Hypothesis Significance Testing (NHST), whose focus is the short-term goal of testing for evidence to support a research hypothesis based on a single trial. In contrast, the HRM focuses on the long-term goal of reliably confirming or disconfirming a research hypothesis via an analysis that combines knowledge from theory and prior empirical evidence together with results from experimental trials.

Second, the model is applied simply, using a binary Bayesian approach that focuses on the model *α*_m_ and *β*_m_ parameters that are routinely calculated for NHST [27,45]. Third, meaningful, general conclusions consistent with the aims of hypothesis testing are drawn to guide research practices, despite uncertainties in the actual values of *R*_0_, *α*, and *β*. Finally, a Monte Carlo simulation shows how the Hypothesis Race Model can be used to cost-effectively find successful treatments for a disease and estimate the total cost for such an endeavor.

## 2. Materials and Methods

### 2.1 Theory

The Hypothesis Race Model builds on Bayes’ theorem, which allows us to update an estimate of the probability that a hypothesis is true, given the results of a test (involving new, nonredundant information) that either supports the hypothesis or its opposite [46]. Bayes’ theorem requires an estimate of this probability prior to incorporating the new information, which for convenience is expressed as the odds ratio, *R*_0_, the probability that the hypothesis is true divided by the probability that it is false. The updated, “posterior” odds ratio, *R*_1_, after applying Bayes’ theorem can be expressed as

*R*_1_ = *R*_0_ * *BF*_1_, where *BF*_1_, the Bayes factor, is given by

*BF*_1_ = (probability of observed results given that the hypothesis is true)/(probability of observed results given that the hypothesis is false).

The results of multiple such tests can be chained together in any order to provide a new estimate of *R* that takes into account all of these test results, together with the prior odds ratio, for example

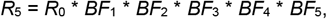

where *BF*_1_-*BF*_5_ represent five independent test results, whose information has not already been included in the estimate *R*_0_.

To demonstrate the principles of the HRM, we begin with an example of testing a repeated measure to determine if an increase has occurred, using a one-tailed paired sample *t*-test (Fig 1). Using the terminology of NHST, we use data from a random sample to test the alternative hypothesis that the mean value of our population has increased, against the null hypothesis that no difference exists. With NHST, empirical evidence never supports the conclusion that the null hypothesis is true [27,35], but for the HRM, evidence is weighed to support the conclusion that the tested hypothesis is true (denoted *H*_T_) or false (denoted *H*_F_); and their estimated probabilities are always related by *P*(*H*_T_) = 1 - *P*(*H*_F_).

**Fig 1.**
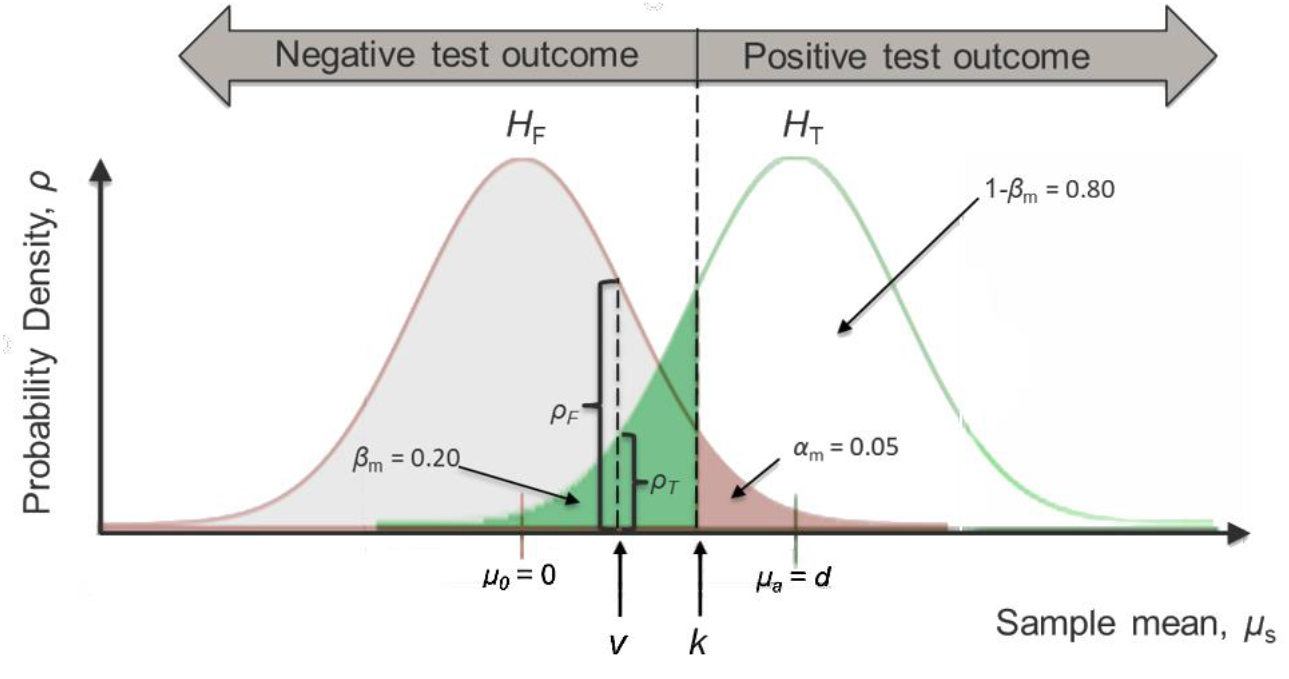
Hypothesis testing: The mean from a sample (*μ*_s_) helps to determine the odds that a hypothesis is true. The model for our example uses a one-tailed, paired sample *t*-test, which assumes that the distribution of sample means will be a Student’s *t*-distribution centered on the value 0 for a false hypothesis (*H*_F_) and *d* for a true hypothesis (*H*_T_). The test is considered positive if *μ*_s_ ≥ *k*, the critical value. For Null Hypothesis Significance Testing (NHST), by convention *k* often represents the point where 5% of the area under the *H*_F_ probability density curve lies to the right of *k* (corresponding to *α*_m_ = 0.05, where *α*_m_ represents the probability of obtaining a positive outcome for a false hypothesis). For the Hypothesis Race Model (HRM) we focus on calculating the Bayes factor (*BF*), the probability that we would get our observed outcome (*μ*_s_) if the hypothesis were true divided by the probability that we would get our observed outcome if the hypothesis were false. For the binary HRM, with *μ*_s_ = *v*, where *v* < *k*, the *BF* for this negative test outcome is given by *β*_m_/1-*α*_m_, where *β*_m_ is the probability of a negative test outcome when the hypothesis is true; and 1 – *α*_m_ is the probability when the hypothesis is false. For the nonbinary HRM, *BF* is given by the ratio *ρ*_T_/*ρ*_F_, where *ρ*_T_ and *ρ*_F_ are the probability densities at point *μ*_s_ = *v* for a true and false hypothesis, respectively.

Our example experiment measures differences between *N* pairs of observations. The observations are assumed to be normally distributed with standard deviation *σ*. The means of the distributions of differences are *μ*_a_ for the alternative hypothesis and *μ*_0_ for the null hypothesis. The raw effect size *μ*_a_ – *μ*_0_ is defined to be *d*\**σ*, where *d* is Cohen’s *d* for the alternative hypothesis. For simplicity, we assume that our random variables have been rescaled such that *σ* = 1 and recentered such that *μ*_0_ = 0, allowing the raw effect size to equal *d*, with *μ*_a_ = *d*.

Fig 1 shows the probability distributions for the means of random samples corresponding to our alternative and null hypotheses: *t* distributions (with *df*=*N*-1, where *N* is the number of pairs in our sample) centered on *μ*_a_ = *d* and *μ*_0_ = 0, respectively. Let *P*(*H*_T_ | prior) be the probability that the alternative hypothesis is true prior to the first experiment and *P*(*H*_F_ | prior) be the probability that the null hypothesis is true (and the alternative hypothesis false) prior to the first experiment. *R*_0_, the prior odds that the alternative hypothesis is true, is given by *P*(*H*_T_ | prior)/ *P*(*H*_F_ | prior). From Bayes’ theorem the posterior odds, after including the new information from the experiment, can be expressed as *R*_1_ = *BF*_1_\**R*_0_, with *BF*_1_ = *P*(*O*_1_ | *H*_T_)/*P*(*O*_1_ | *H*_F_), where *O*_1_ is the test-statistic outcome of our first experimental trial.

For our binary HRM, *O*_*1*_ takes only two possible values, positive or negative, which depend upon whether *v*_1_, the sample mean outcome for our first trial, exceeds or equals a threshold, *k*, the same critical value that defines a positive test result for NHST. For a positive result, *BF*_+_ = *P*(+ | *H*_T_)/*P*(+ | *H*_F_) = 1-*β*/*α*, and for a negative result *BF*_-_ = *P*(-| *H*_T_)/*P*(-| *H*_F_) = *β*/1-*α*; where *α* is the probability of a type I error (rejecting the null hypothesis when it is true) and *β* the probability of a type II error (accepting the null hypothesis when it is false).

For our nonbinary HRM, *O*_1_ represents the outcome *v*_1_, and *BF*_1_ = *ρ*_T1_/*ρ*_F1_, where *ρ*_T1_ and *ρ*_F1_ are the probability densities at point *v*_1_ when the alternative hypothesis or null hypothesis is true, respectively. We refer to this model as our nonbinary model because *BF*_1_ varies continuously as a function of *v*_1_.

We focus primarily on the binary HRM because its simplicity facilitates evaluation of the effects of bias on our estimates of research progress toward confirmation/disconfirmation of our alternative hypothesis. It also corresponds closely to NHST with power analysis, so it builds on concepts that are familiar to the research community [42,45]. An important difference is that NHST typically does not account for the effects of bias on the actual values *α* and *β*, where model values are used instead, *α*_m_ and *β*_m_ (Fig 1), calculated from the statistical testing model. For the HRM, bias terms are included, *α* = *α*_m_ + *α*_b_ and *β* = *β*_m_ + *β*_b_, where *α*_b_ and *β*_b_ represent the amounts by which the actual values deviate from the model values. Such deviations can be caused by various forms of bias, including selection bias, placebo effect, observer-expectancy effect, publication bias, methodological errors, etc. For example, Simmons et al. (2011) [47] showed with thousands of computer-generated random samples that allowing flexibility in choosing study endpoints, sample size, and typical methods of data analysis lead to finding *p* ≤ 0.05, 61% of the time, corresponding to *α*_m_ = 0.05 and *α* = 0.61.

Fig 2 provides an illustration of how to conceptualize the HRM, showing three scenarios beginning with an assumption of a prior *R*-value and following a series of positive outcomes, ultimately leading to reaching the “finish line,” the point where we are confident enough that our alternative hypothesis is true that we are ready to take action. Examples of such actions could be warning the public concerning a health hazard; marketing a product; or allocating resources for further testing, with the goal of further increasing our confidence in the tested hypothesis.

**Fig 2.**
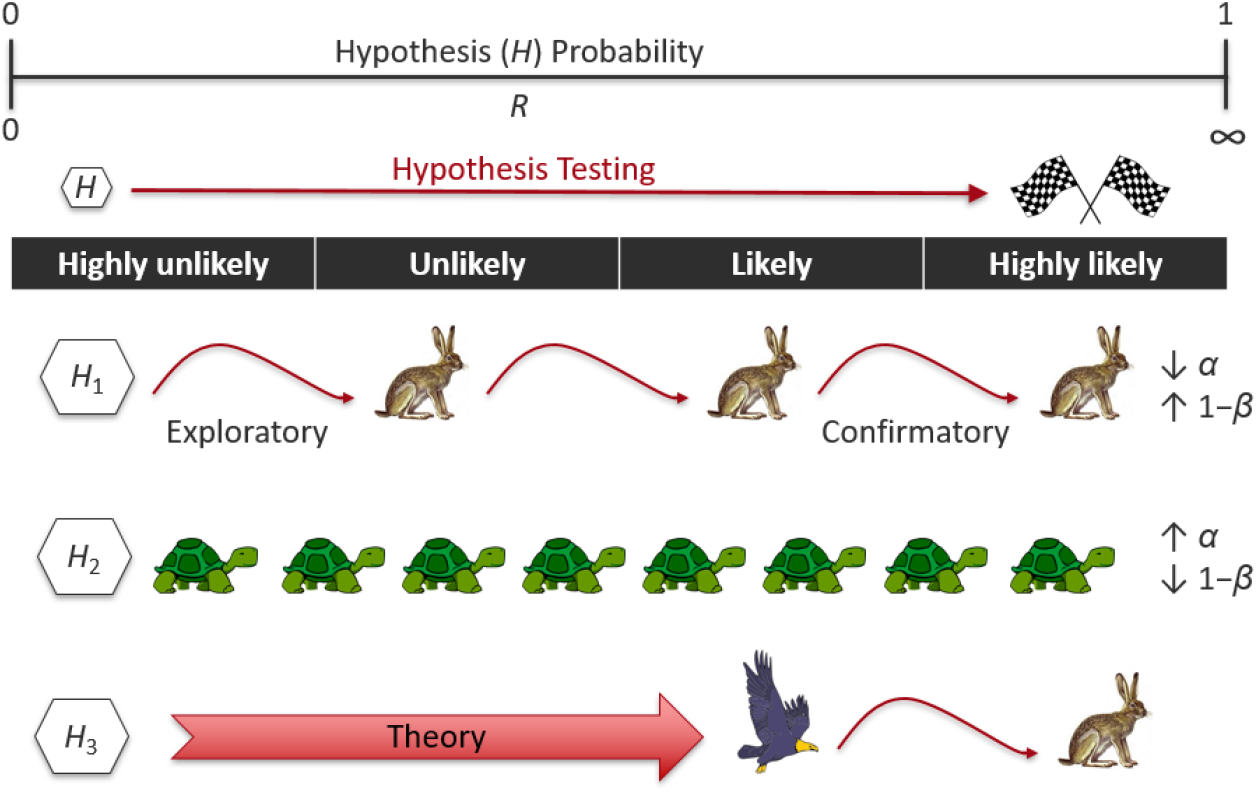
The Hypothesis Race Model. A research hypothesis is deemed confirmed when the combined weight of evidence from experimental trials moves the probability (or odds, *R*-value) that the hypothesis is true across the “finish line,” the point where we consider the hypothesis likely enough to warrant further action. Illustrated here are three paths toward confirmation, beginning with hypotheses (*H*_1_-*H*_3_) considered highly unlikely prior to empirical testing and consideration of theory. For *H*_1_ the alpha level (*α*) is low and power (1-*β*) high enough that each positive test outcome from experimental trials results in great strides toward confirmation. Conversely, high *α* and low 1-*β* slows progress toward confirmation (*H*_2_). For *H*_3_, available theory permits a big head start in the race toward confirmation.

Negative test results reduce the net *R*-value (Fig 3), and can even lead to disconfirmation of a hypothesis, the point where we can be confident enough that the tested hypothesis is false, that no further testing is warranted. The *R*-value after *n* experiments can be expressed as the product of the Bayes factors times the prior odds, *R*_n_ = *BF*_1_\**BF*_2_\**BF*_3_ … \**BF*_n_\**R*_0_. Expressing the HRM using a log_10_ scale simplifies the arithmetic, leading to confirmation distance *D*_c_ = log(*R*_n_) – log(*R*_0_) = log(*BF*_1_) + log(*BF*_2_) + log(*BF*_3_) … + log(*BF*_n_). The right-hand side of this equation is independent of our assumptions for *R*_0_ and can be used as a measure of the progress of our research toward confirmation or disconfirmation of the tested hypothesis.

**Fig 3.**
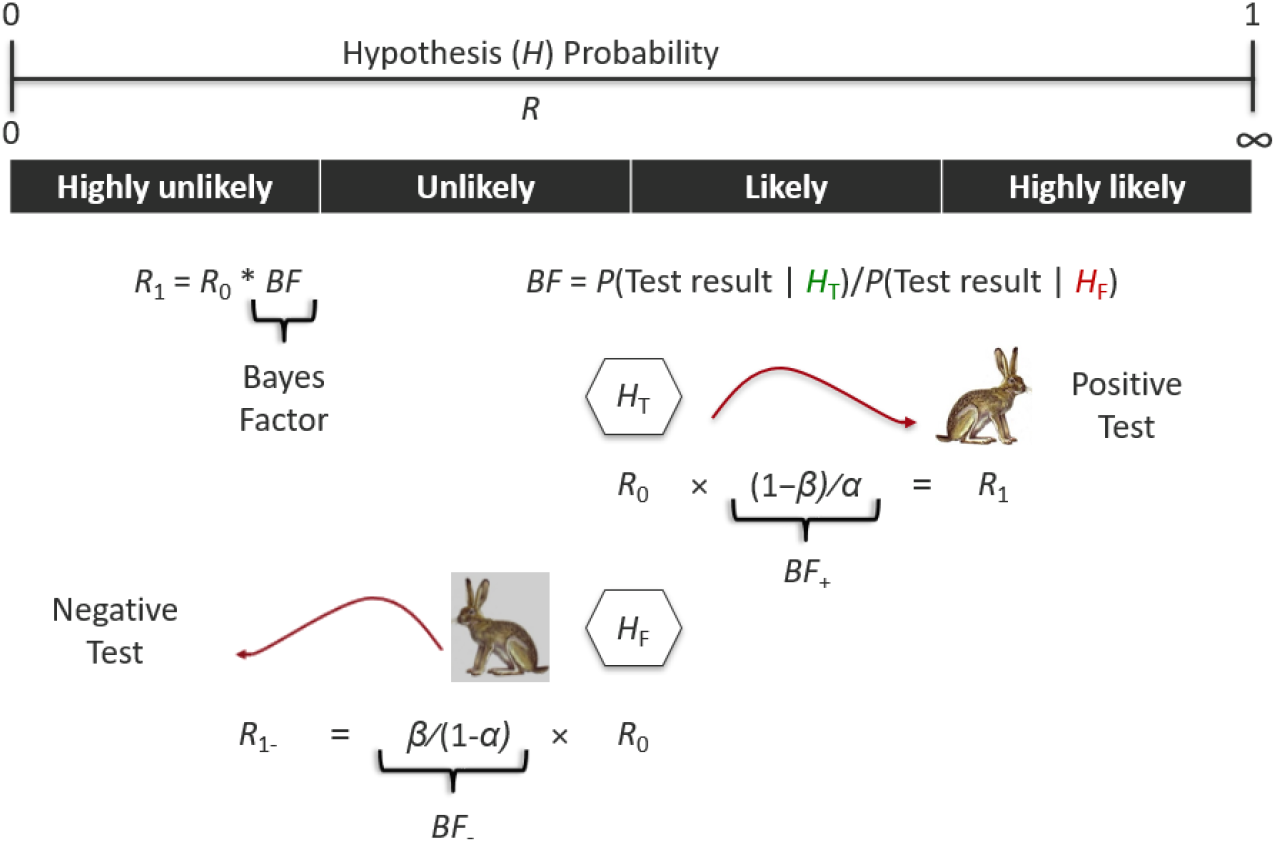
The Hypothesis Race Model (HRM) allows both confirmation and disconfirmation of hypotheses. Positive test results increase *R*, while negative results decrease it. Chance affects test outcomes, but on average, after multiple trials, true hypotheses (*H*_T_) will tend to move right toward confirmation; and false hypotheses (*H*_F_), left toward disconfirmation. This movement is key to the separation of true from false hypotheses through empirical testing. However, the actual values of *α* and *β* are unknown. This article considers the utility of estimating *α* and *β* to facilitate the evaluation of research progress toward confirmation or disconfirmation.

For the binary HRM, only two values of *BF* are possible, *BF*_+_ and *BF*_-_, so *D*_c_ is determined by the frequency of positive (*f*_*+*_) and negative (*f*_-_) outcomes, *D*_c_ = *f*_+_*log(*BF*_+_) + *f*_-_*log(*BF*_-_). If we define *V*_c_ as the average confirmation distance traveled per trial, equal to the expected value of *D*_c_ divided by the number of trials, *n, V*_c_ becomes a measure of the efficiency of research toward the goal of confirming or disconfirming our research hypothesis. *V*_c_ = E(*D*_c_/*n*) = *w*_+_*log(*BF*_+_) + *w*_-_* log(*BF*_+_), where *w*_+_ = 1-*β* and *w*_-_ = *β* for true hypotheses (*d*>0); and *w*_+_ = *α* and *w*_-_ = 1-*α* for false hypotheses (*d*=0). Thus, for true hypotheses, *V*_c_ = *V*_c_(*H*_T_) = (1-*β*)*log((1-*β*)/*α*) + *β*\*log(*β*/(1-*α*)); and for false hypotheses *V*_c_ = *V*_c_(*H*_F_) = *α*\*log((1-*β*)/*α*) + (1-*α*)*log(*β*/(1-*α*)). Actions that increase the magnitude of *V*_c_, increasing *V*_c_ for true hypotheses and decreasing it for false hypotheses, improve the cost-effectiveness of research.

The problem with these equations is that the actual values of *α* and *β* are unknown. With excellent control of all forms of bias, the model value, *α*_m_, can become a good approximation to *α*; however, reliably finding a good model approximation for *β* is generally impossible, because *β* and *β*_m_ are directly related to the effect size, *d*, for our experiment. If we knew the value of *d*, there would obviously be no point in hypothesis testing.

However, through judicious estimates for the values of *α* and *d* it is possible to apply the HRM to evaluate research progress, confirm or disconfirm tested hypotheses, and promote more efficient use of research resources. We begin by exploring the effect of estimates on the value of *V*_*c*_, denoted by the subscript *e*: 1) if the tested hypothesis is true,

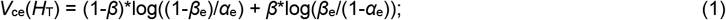

and 2) if the tested hypothesis is false,

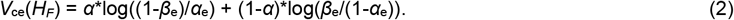

In practice, the estimated confirmation distance traversed in the course of repeated, identical experiments, would be given by

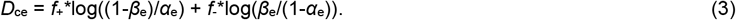

The actual confirmation distance is a product of chance, but if we can ensure that *V*_ce_(*H*_T_)>0 and *V*_ce_(*H*_F_)<0, then with a large number of trials, *D*_ce_ should tend toward confirmation of true hypotheses and disconfirmation of false ones.

The appendix shows that *V*_ce_ is always positive for true hypotheses provided that we choose conservatively low estimates for *d*_e_ and hence power, 1-*β*_e_; and *V*_ce_ is always negative for false hypotheses for conservatively high estimates for *α*_e_. *V*_ce_(*H*_T_) is maximized when 1-*β*_e_ = 1-*β*, and (substantially) higher values of 1-*β*_e_ can decrease *V*_ce_(*H*_T_) to become < 0, allowing disconfirmation of a true hypothesis, with a sufficiently large number of trials. Conversely, *V*_ce_(*H*_F_) is minimized when *α*_e_ = *α*, and lower values of α_e_ can increase *V*_ce_(*H*_F_) to become > 0, allowing confirmation of a false hypothesis.

Thus, the HRM can be adjusted for anticipated bias to prevent confirmation of false hypotheses by choosing a sufficiently large value of *α*_e_ to ensure that it is no smaller than *α*. For example, if we knew that with *α*_m_ = 0.05 we could anticipate positive test results in 61% of cases (as with Simmons et al.’s example [47]), then by setting *α*_e_ = 0.61, we could still separate true from false hypotheses, albeit at a much slower rate per trial than if there had been no bias.

Fig 4 illustrates that when *d*_e_ >> *d* > 0, then *V*_ce_(*H*_T_) becomes negative, which would violate our requirement that *V*_ce_ be positive for a true hypothesis. In contrast, when underestimating *d* in the absence of bias, *V*_ce_ is always positive for true hypotheses and negative for false ones. However, if 0 < *d*_e_ << *d*, then *V*_ce_ would become much smaller than *V*_c_, resulting in an underestimation of actual research progress from our experiments. In practice, very small effect sizes may be of no interest, so a reasonable strategy would be to choose a conservatively low value of *d*_e_ (and 1-*β*_e_), but large enough to capture effect sizes of interest.

**Fig 4.**
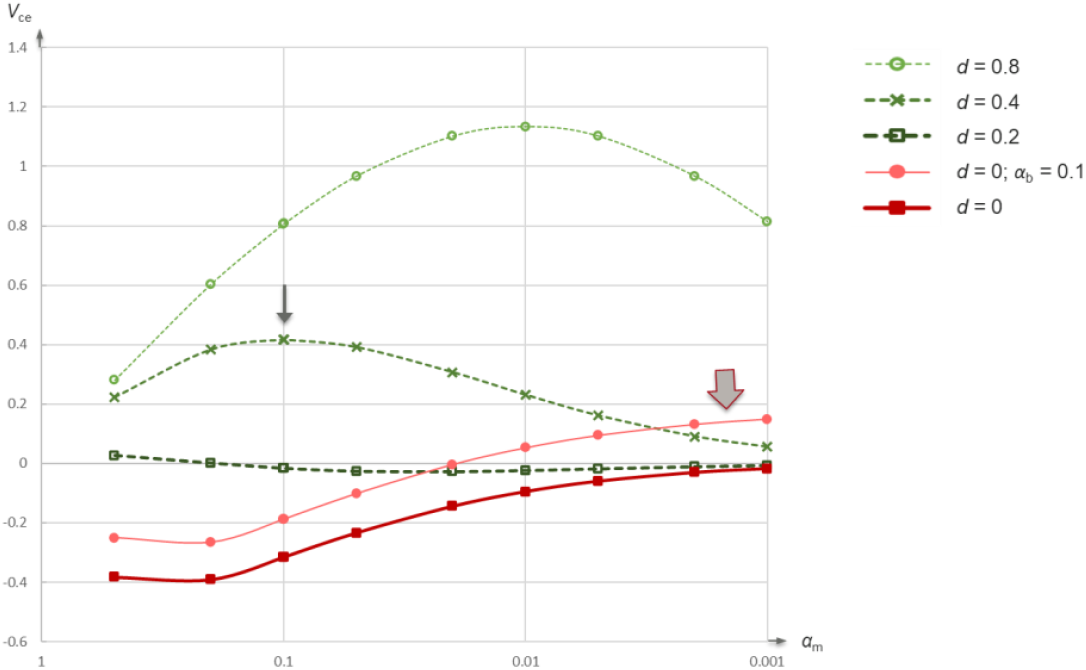
Mean confirmation velocity estimate, *V*_ce_, plotted against *α*_m_, the statistical model’s *α*-level, for sample size *N* = 20, using effect size estimate *d*_e_ = 0.4. Each line represents the graph of *V*_ce_ for actual effect sizes from *d* = 0 to *d* = 0.8, and one case with bias *α*_b_ = 0.1. For *d* = 0.4, maximum *V*_ce_ is achieved by choosing *α*_m_ at the peak of the curve (small arrow). The higher *d* = 0.8 allows higher *V*_ce_, but the lower *d* = 0.2 < *d*_e_ can result in negative *V*_ce_ and incorrect disconfirmation of a true hypothesis. For false hypotheses (*H*_F_: *d* = 0), if our estimate assumes no bias (i.e. *α*_e_ = *α*_m_), then even small amounts of actual bias (*α*_b_ = *α* – *α*_m_) can cause *V*_ce_ > 0 for small values of *α*_m_ (large arrow), allowing incorrect confirmation of a false hypothesis.

Fig 4 also demonstrates the importance of choosing an optimal value of *α*_m_ (for now considered with *α*_b_=0) to maximize *V*_ce_(*H*_*T*_). Provided that *d*_e_ ≤ *d*, the plots of *V*_ce_(*H*_T_) vs. *α*_m_ have a single maximum value > 0. This optimal value of *α*_m_ could be higher or lower than the conventional 0.05, depending upon the values of *N, d*_e_ and *d*. In addition, small amounts of added bias, if not included in our estimate *α*_e_, could result in a positive *V*_ce_(*H*_F_) (Fig 4, large arrow).

For some forms of bias, such as publication bias, *α* would be expected to be proportional to *α*_m_ [15]. However, bias due to methodological errors could be independent of *α*_m_ and thus *α*_b_ could become proportionally much larger in relation to *α*_m_ as *α*_m_ becomes very small. A methodological error that changes a positive test result to negative in 1% of cases would normally have little effect on *V*_c_(*H*_*T*_) or *V*_c_(*H*_F_) because the error would change little the actual power, 1-*β* (e.g., for a well-powered study, change from 0.80 to approximately 0.79), or the *α*-level (e.g., if *α*_m_ = 0.001, then *α* would become ∼ 0.00099). However, a methodological error that changes a negative test result to positive in 1% of cases would have a dramatic effect on *V*_c_ for small *α*_m_. For example, for *α*_m_ = 0.001, *α* would become ∼ 0.011.

To err is human, so even if we were to consider negligible the risk of *p*-hacking or deliberate data tampering, which can be considered part of the bias due to “methodological errors,” this form of bias can never be zero. Therefore, it makes sense to include at least a small amount of anticipated bias in our estimate, *α*_e_, to reduce or eliminate the risk of incorrectly confirming a false hypothesis. Doing so impacts the optimal choice of sample size, *N*, as illustrated by Table 1: In the absence of bias, larger studies are slightly more efficient in terms of *V*_c_(*H*_T_), and hence *V*_ce_(*H*_T_), per participant. However, if we model methodological errors with *α*_e_ ≈ *α* = *α*_m_ + *α*_b_, where *ε*_M_ is the risk of changing a negative outcome to a positive one, then *α*_b_ = *ε*_M_ * (1-*α*_m_). For *ε*_M_ = 0.001 (arguably an unrealistically small estimate) or 0.01, Table 1 shows that *V*_c_(*H*_T_) per study participant in both cases is optimized for *N* = 20 or 80 compared with *N* = 320.

**Table 1.**
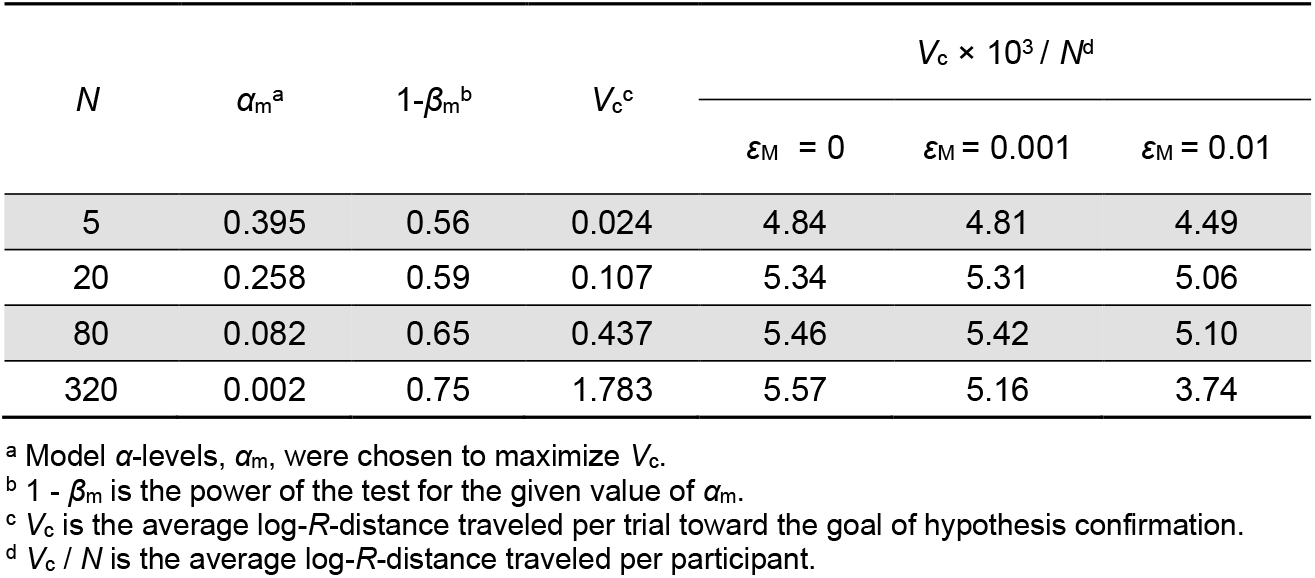
*V*_c_ vs. sample size (*N*) for the binary HRM, with effect size *d* = 0.2. Larger sample sizes become less cost-effective when the risk of bias from methodological errors (*ε*_M_) is considered with *α* = *α*_m_ + *α*_b_, where *α*_b_ = *ε*_M_ * (1-*α*_m_). Estimates for *α* and *d* are assumed equal to actual values for this example.

| $N$ | $\alpha_m^a$ | $1-\beta_m^b$ | $V_c^c$ | $V_c \times 10^3 / N^d$ | | |
| --- | --- | --- | --- | --- | --- | --- |
| | | | | $\epsilon_M = 0$ | $\epsilon_M = 0.001$ | $\epsilon_M = 0.01$ |
| 5 | 0.395 | 0.56 | 0.024 | 4.84 | 4.81 | 4.49 |
| 20 | 0.258 | 0.59 | 0.107 | 5.34 | 5.31 | 5.06 |
| 80 | 0.082 | 0.65 | 0.437 | 5.46 | 5.42 | 5.10 |
| 320 | 0.002 | 0.75 | 1.783 | 5.57 | 5.16 | 3.74 |

<sup>a</sup> Model $\alpha$ -levels, $\alpha_m$ , were chosen to maximize $V_c$ .
<sup>b</sup> $1 - \beta_m$ is the power of the test for the given value of $\alpha_m$ .
<sup>c</sup> $V_c$ is the average log- $R$ -distance traveled per trial toward the goal of hypothesis confirmation.
<sup>d</sup> $V_c / N$ is the average log- $R$ -distance traveled per participant.

So if a study’s cost is proportional to the number of participants, a more efficient utilization of research resources could be obtained by breaking larger trials (*N*>100) into multiple, mid-size trials (*N*<100), provided that the risk of methodological errors in each trial would be independent of the risk in the others. Separate research teams for each trial might help to promote such statistical independence. Although larger sample sizes and smaller *α*_m_ may promote replicability of research results [48], the opposite--smaller sample sizes and larger *α*_m_—could increase the overall cost-effectiveness of *confirming* research hypotheses, by lessening the impact of methodological errors. The next section describes another situation where smaller sample sizes conserve research resources.

### 2.2 Horse Race Variant of the HRM

The Horse Race Variant describes the special situation where many hypotheses are considered, each with a small chance of being a “winner” (for example, when testing multiple chemotherapy compounds to see which would be effective in thwarting tumor growth), but where we are only interested in finding one or a few winners in the pack. In this situation, research resources (e.g., number of petri dishes processed) might be conserved by not systematically testing every hypothesis. Small trials could be utilized to identify promising candidates, where research resources would then be concentrated.

To illustrate one such approach, Monte Carlo simulations were employed, for both the binary and nonbinary applications of the HRM, starting with a database of 1,000 candidate hypotheses, one of which was a “winner,” with actual *d*=0.2 or *d*=0.4. The remaining hypotheses had *d*=0. The process was to test one hypothesis at a time and reevaluate *D*_ce_ after each trial, stopping testing when a hypothesis had traversed the distance from log(*R*) = -3 to +2, or had become less likely than the remaining hypotheses. In other words, we began with an assumption that the probability of a true hypothesis started at ∼0.001 and were satisfied when this probability was determined to be at least ∼0.99. We tested trial sizes of *N* = 5, 10, 20, 40, 80, and 160 with an estimated effect size *d*_e_ = 0.2 and assumed *α*_b_ = 0. For the binary application of the HRM, we chose the value of *α*_m_ that maximized *V*_ce_(*H*_T_) for the chosen sample size; and for the non-binary application the *α*-level was unnecessary. This process continued until a winning hypothesis was found, not necessarily the sought true hypothesis. For statistical comparisons, 5000 iterations were performed for each combination of *N, d*, and binary/nonbinary applications. Mean values for number of participants needed (i.e., product of sample size and mean number of trials) were compared using two-tailed, two-sample *t*-tests, considering *p* < 0.001, uncorrected, to be statistically significant. For each of the 4 groups (binary/nonbinary × 2 effect sizes), the sample size requiring the lowest total number of participants was used as the reference for comparison with the remaining 5 sample sizes, for a total of 20 comparisons.

## 3. Results

For the Horse Race Variant, in each of the four groups the lowest total number of participants was achieved with sample sizes *N* = 10 or 20, which were significantly more cost-effective in terms of total number of participants than sample sizes N = 80 or 160 (Fig 5). Despite the relatively small standard errors for mean number of participants, the ranges for total number of participants spanned from many times smaller (∼10-20) to many times greater (∼5-10) than the means, due to chance. The percentage incorrectly identified winners for each set of 5000 iterations ranged from 0.06% to 0.56%, consistent with expectations, given that each winner required a log(*R*) value of at least 2.

**Fig 5.**
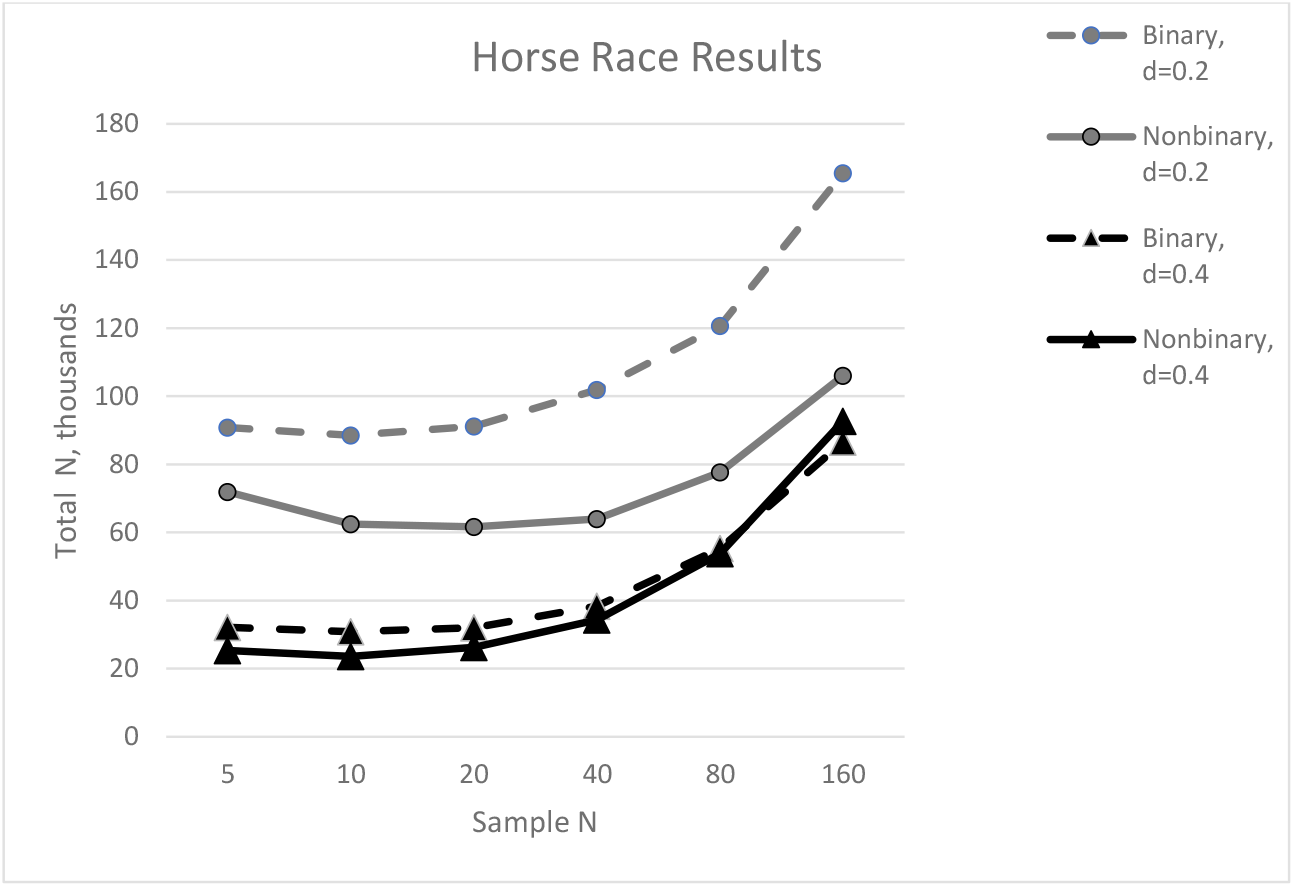
Results of the Horse Race Variant, showing the average total cost in terms of number of study participants (Total *N*) needed to repeatedly test (using fixed sample size, Sample *N*) to find the winning hypothesis from a field of 1,000 hypotheses, each with a 1/1000 chance of being a winner. This total cost was lowest for trials having only 10 or 20 participants compared with larger trial sample sizes.

## 4. Discussion

The HRM fixes long-standing problems with the NHST model that often lead researchers to draw unrealistic conclusions when designing research studies and evaluating their results. Compared with other Bayesian models the HRM is easily adopted by researchers familiar with the NHST model and concepts such as *α*-level and power analysis. The *R*-value might present new conceptual challenges for some researchers, but it lies at the core of the paradigm shift required for proper interpretation of research results: Each new result should be considered supporting evidence, which together with theory and previous empirical results leads to an updated *R*-value estimate and corresponding probability that a hypothesis is true. The goal of empirical testing is not to determine the “truth” from a single study, but to help determine if the weight of collected evidence motivates a shift in existing conclusions and corresponding actions.

Adoption of the HRM should benefit many fledgling researchers because it provides an intuitive framework that circumvents pitfalls associated with the NHST paradigm. For example, using the standard NHST approach it is natural to conclude that individual studies showing statistically significant results should lend significant credibility to the tested hypotheses [49]. In the author’s experience, papers submitted for publication from inexperienced colleagues often read something like “Authors A have proven B and now we have proven C.” However, as experienced researchers understand, preliminary research findings “are in need of independent replication” [6,22,50–52]. Points readily understood from the HRM perspective are that a single study result alone should not lead to fixed conclusions; and that hypothesis “confirmation” requires the inclusion of results from all empirical tests of the hypothesis, including negative results, which are often suppressed due to publication bias. Another point readily understood is that a small, marginally statistically significant study showing a large effect size is of limited value toward confirmation of the tested hypothesis [53], in consideration of *R*- and *β*-values; while from the NHST perspective such a result might seem more promising than a larger study with the same *p*-value.

Sometimes the NHST paradigm can mislead experienced researchers [13,54]. The NHST directs the focus of attention to the statistical significance of a single research study (or meta-analysis as a proxy for a single, larger study); but the ultimate goal should be confirmation of the tested research hypothesis. Other factors deserving consideration include the *R*-value, sources of bias affecting the *α*-level, and the degree to which the tested hypothesis aligns with the clinically relevant hypothesis of interest. Hypotheses that from the outset seem highly improbable or are selected from huge “data-mining” operations should not be granted the same credence as hypotheses that are well supported by theory and/or previous empirical results.

Conclusions may differ depending upon the paradigm used, as exemplified by the recent controversy over the optimal *α*-level for statistical significance of novel research findings [2,48,55–57]. From the NHST perspective, lowering the α-level from 0.05 to 0.005 would be beneficial by lowering the rate of false positive results; and larger sample sizes could mitigate publication bias [52] by including a greater share of existing test results. However, in some cases this change in *α*-level could increase the overall research cost for confirmation or disconfirmation of tested hypotheses. For example, instead of using one large, well-powered sample, smaller samples from independent research groups could serve as a hedge against small risks of methodological error and other potential sources of bias, thereby lowering total costs for hypothesis confirmation. And when testing many medicines against illness simultaneously (Horse Race Variant), smaller sample sizes with larger *α*-levels can lower the overall cost of finding an effective medication.

Conclusions also differ concerning the value of positive studies later shown not to be replicated. Researchers from Amgen noted that only 6/53 of their studies confirmed previous findings, a low success rate described as “not sustainable” [58]. Some have suggested that researchers and journals publishing false-positive studies should be held accountable for publishing such “problematic data” [59]. The NHST perspective understandably promotes such views, but a broader, more realistic view is possible with the HRM. For example, if Amgen’s 6 of 53 replicable studies represented a narrowing of the field from 1000 candidate drugs to only 53 drugs and finally 6 effective drugs, then the process could be viewed as an overall success, despite the occurrence of false positives. Variants of the HRM could help to maximize the efficiency of the process, which can prove expensive if candidate drugs all begin with a low probability of success.

As a case in point, consider the hydroxychloroquine controversy during the COVID-19 epidemic. Although large studies of high quality did not confirm the broad hypothesis that hydroxychloroquine could successfully treat COVID-19 [60,61], at the time of publication of their article (since retracted due to methodological concerns [62]), such evidence was lacking when researchers published results from a small open-label study with statistical significance of *p* < 0.001 testing the specific hypothesis that COVID-19 virus would less often be detected in patients after treatment with hydroxychloroquine [63]. Given a similar situation, possibly during a future pandemic, what course of action should researchers recommend to policy makers? Although parameters for the HRM cannot be known exactly (including *R*_0_, *α*-level, *β*, and target *R*-value at which specific actions should be recommended; as well as relevance of the tested hypothesis, methods, and population), the HRM provides a common framework for a more nuanced discussion of relevant factors. Most important is to recognize the uncertainty inherent in such judgment calls and inform the design of studies to move the collected results closer to confirmation or disconfirmation so that appropriate actions are taken.

As demonstrated by the nonbinary HRM in the current paper, multiple Bayesian approaches are possible that can in some cases provide more accurate and/or cost-effective means of designing studies and evaluating research results. However, the current study focuses on the dichotomous approach by Ioannidis (2005) [15], classifying findings as positive or negative, because of the greater simplicity and clarity provided by this approach when estimating and including factors in the model such as bias. Given the uncertainty inherent in model parameters, clarity may be preferable over attempts at greater precision, though researchers should tailor their approaches according to their individual needs.

A word of caution concerning recently popularized Bayesian approaches to analyzing research data to determine statistical significance of individual trials [46,64]: These sophisticated Bayesian approaches, like NHST, focus on individual results rather than the overall path toward ultimate hypothesis confirmation [2,65]. They may effectively lower the nominal *α*-level to less than 0.05 [34], potentially increasing research costs.

Limitations of the current study include the lack of preregistration of study hypotheses and independent testing of methods used. Both of these shortcomings are addressed by making the software and data for this study publicly available so that other researchers can confirm the validity of this work and/or provide corrections. The author has also reproduced this work, following the described methods.

## 5. Conclusions

Despite repeated calls for a Bayesian approach to hypothesis testing in medicine and the social sciences, NHST remains the most popular approach [3,42,66]. The HRM facilitates the addition of a Bayesian approach, building upon familiar NHST-concepts. It is an intuitively appealing framework that provides a common language for discussion as well as the foundation for mathematical models that can help to reduce the cost of empirical hypothesis testing.

## Supplementary Materials

Two English-language video presentations on the HRM by R.E.K.J. are available through the Swedish Psychiatric Association’s website at https://www.svenskapsykiatrikongressen.se, with links also available at www.robertkelly.us/presentations.

## Data Availability Statement

The original data and R code for the current study are openly available in the GitHub repository at github.com/robertkellymd/hrm.

## Acknowledgments

Many thanks for valuable feedback from Jonas Eberhard, Lund University; Matthew J. Hoptman, Nathan S. Klein Institute and New York University; and Hugo Eberhard, University of Cambridge.

## Abbreviations

The following abbreviations are used in this manuscript:

BF: Bayes factor
HRM: Hypothesis Race Model
NHST: Null Hypothesis Significance Testing

## Appendix

This appendix elucidates the conditions necessary to estimate research progress and distinguish true from false hypotheses to any desired degree of certainty, through repeated empirical testing.

The estimated log-distance traversed toward the goal of confirmation or disconfirmation of a study hypothesis is given by Equation 3, *D*_ce_ = *f*_+_*log((1-*β*_e_)/*α*_e_) + *f*_-_*log(*β*_e_/(1-*α*_e_)), where *f*_+_ and *f*_-_ are the frequencies of positive and negative test outcomes, *α*_e_ is the estimated type I error, and *β*_e_, the estimated type II error. The estimated power of the test is 1-*β*_e_, which corresponds directly to the estimated effect size, *d*_e_, for a true hypothesis.

Our estimate for *D*_ce_ can allow us to distinguish true from false hypotheses provided that its expected value is positive for true hypotheses and negative for false hypotheses, or equivalently, from Equations 1 and 2, that *V*_ce_(*H*_T_) = (1-*β*)*log((1-*β*_e_)/*α*_e_) + *β*\*log(*β*_e_/(1-*α*_e_)) > 0 and *V*_ce_(*H*_F_) = *α*\*log((1-*β*_e_)/*α*_e_) + (1-*α*)*log(*β*_e_/(1-*α*_e_)) < 0; where *V*_ce_ = E(*D*_ce_/*n*), *n* is the number of trials, *α* is the actual type I error, and *β*, the actual type II error. The actual power of the test, 1-*β*, corresponds directly to the actual effect size, *d*, for a true hypothesis (Fig 1).

For simplicity we assume that all tests are identical, but differ in outcome, positive or negative, due to chance. To ensure that our empirical tests serve their fundamental purpose, we also assume that 1-*β* > *α* (equivalent to *β* < 1-*α*); and correspondingly, that 1-*β*_e_ > *α*_e_.

We demonstrate the following six properties of *V*_*ce*_, then summarize their significance for the process of distinguishing true from false hypotheses via empirical testing.

> P1: When *β*_e_ = *β*, corresponding to *d*_e_ = *d*, then *V*_ce_(*H*_T_) is maximized.
>
> P2: When *β*_e_ ≥ *β*, corresponding to *d*_e_ ≤ *d*, then *V*_ce_(*H*_T_) > 0.
>
> P3: When *β*_e_ < *β*_0_, then *V*_ce_(*H*_T_) < 0 for some value *β*_0_ that lies between 0 and *β*, which corresponds to a value *d*_0_ such that *d*_e_ > *d*_0_ > *d*.
>
> P4: When *α*_e_ = *α*, then *V*_ce_(*H*_F_) is minimized.
>
> P5: When *α*_e_ ≥ *α*, then *V*_ce_(*H*_F_) < 0.
>
> P6: When *α*_e_ < *α*_0_, then *V*_ce_(*H*_F_) > 0 for some value *α*_0_ that lies between 0 and *α*.

Proof of P1: From Equation 1 we have

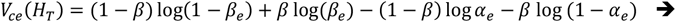

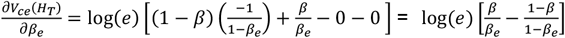, which means that

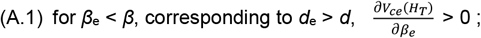

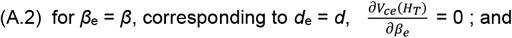

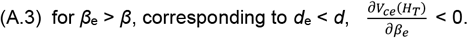

Thus, V_ce_(*H*_T_) reaches its maximum value when *β*_e_ = *β*.

Proof of P2: We show that as *β*_*e*_ → (1 − *α*_*e*_), *V*_*ce*_ (*H*_*T*_) → 0. From Equation 1 we have

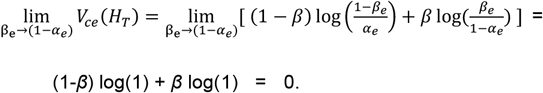

Together with A.3, this means that V_ce_(*H*_T_) > 0 for *β*_e_ on the interval [*β*, 1-*α*_e_).

Proof of P3: We show that as *β*_*e*_ → 0, *V*_*ce*_ (*H*_*T*_) → −∞. From Equation 1 we have

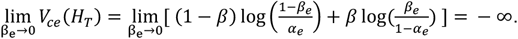

This means that as *β*_e_ becomes successively smaller, at some point V_ce_(*H*_T_) will become < 0.

Proof of P4: From Equation 2 we have

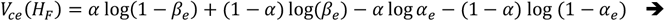

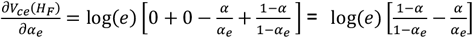, which means that

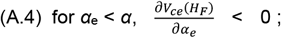

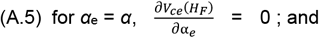

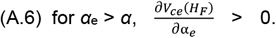

Thus, V_ce_(*H*_F_) reaches its minimum value when *α*_e_ = *α*.

Proof of P5: We show that as *α*_*e*_ → (1 − *β*_*e*_), *V*_*ce*_ (*H*_*F*_) → 0. From Equation 2 we have

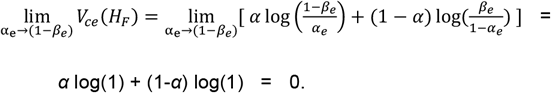

Together with A.6, this means that V_ce_(*H*_*F*_) < 0 for *α*_e_ on the interval [α, 1-*β*_e_).

Proof of P6: We show that as *α*_*e*_ → 0, *V*_*ce*_ (*H*_*F*_) → +∞. From Equation 2 we have

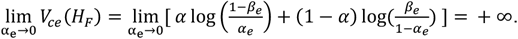

This means that as *α*_e_ becomes successively smaller, at some point *V*_ce_(*H*_F_) will become > 0.

In summary, we can ensure that our estimate *D*_ce_ allows separation of true from false hypotheses, provided that we use conservatively small estimates for effect size, *d*_e_, and conservatively large estimates for *α*-level, *α*_e_. When *d*_e_ >> *d*, true hypotheses may be disconfirmed, and when *α*_e_ << *α*, false hypotheses may be confirmed with repeated, unlimited empirical testing. However, choosing estimates of *d*_e_ << *d* or *α*_*e*_ >> *α* will result in underestimating the actual progress toward confirmation or disconfirmation. Choosing estimates as close as possible to their actual values will provide the most accurate estimates of *V*_c_ and *D*_c_.

Although the actual values of *d* and *α* generally cannot be known, it is possible to make decisions based on a series of what-if scenarios; and as discussed in the body of the text, in practice it is often possible to make well-informed decisions based on reasonable assumptions concerning the possible values of *d* and *α*.

